# A knowledge-based distance metric highlights underperformance of variant effect predictors on gain-of-function missense variants

**DOI:** 10.1101/2025.07.23.666325

**Authors:** Lukas Gerasimavicius, Joseph A. Marsh

## Abstract

Most current variant effect predictors (VEPs) are less effective at identifying non-loss-of-function (non-LOF) disease variants, which arise due to a gain of function (GOF) or dominant-negative (DN) mechanism. Previously, we showed that Extent of Disease Clustering (EDC), a structure-derived metric, can be used to prioritize putative non-LOF genes due to the tendency of their pathogenic missense variants to cluster. We also introduced a structure-derived spatial distance metric, Spatial Proximity to Disease Variants (SPDV), which proved useful for scoring RyR1 variants. Here, we apply EDC and SPDV across thousands of human disease genes. In comparison with 72 variant effect predictors (VEPs), SPDV exceeds or is at least as effective as half of the tested models at identifying dominant disease variants. When applied to GOF variants, SPDV performs similarly to top VEPs. We show that SPDV performance improves with increasing target EDC values and identify an EDC threshold after which SPDV consistently surpasses all other predictors. Its use as an analytical metric enables the identification of a subset of putative GOF disease genes where current VEP methods underperform relative to SPDV, indicating they may be improved by incorporating spatial distance information.

## Introduction

With the ever-growing abundance of population sequencing data, computational approaches have become essential in assessing the clinical relevance of newly discovered genetic variants and prioritizing them for in-depth characterization. The development of numerous *in silico* variant effect predictors (VEPs) has been fueled by the availability of large-scale variant databases^1^. These tools primarily focus on evaluating missense mutations, which alter the identity of a single amino acid residue and can affect protein activity, stability, and intermolecular interactions^2–4^.

Complementing these computational efforts, experimental methods, such as deep mutational scanning (DMS), have enabled functional interrogation of all possible single amino acid substitutions in a gene for a given phenotype, using competitive assay experiments, with hundreds of targets now explored or currently under investigation^5–8^. Despite its potential, DMS remains constrained at scale by high resource demands and the complexity that arises when attempting to accurately model variant effects from the perspective of multiple underlying functions or phenotypes. While DMS may eventually provide comprehensive functional maps across the proteome, computational predictions remain indispensable for scalable variant interpretation^9–11^.

However, current VEPs – whether based on evolutionary conservation, physicochemical features, or more recent machine learning approaches and language models – consistently underperform in predicting the effects of non-loss-of-function (non-LOF) disease variants^12–14^. Loss-of-function (LOF) mutations, the most thoroughly characterized mechanism, cause disease phenotypes by destabilizing protein structures or disrupting essential interactions, particularly visible in constrained haploinsufficient genes. However, while loss-of-function is the most well understood and commonly assumed molecular mechanism, disease also manifests through alternative mechanisms, such as gain-of-function (GOF) and dominant-negative (DN) effects, which contribute significantly to clinical disease, yet are less well captured by existing predictive frameworks^14–16^. GOF mutations may induce constitutive activity or confer entirely new functions, whereas DN variants impair wild-type protein function through direct competition or incorporation into dysfunctional complexes. Although these non-LOF variants often exert milder effects on protein stability, they frequently result in severe clinical phenotypes—an inconsistency not adequately reflected in conservation-based scoring systems^17–20^. Indeed, even unsupervised or conservation-informed methods fail to generalize across GOF and DN pathogenic mechanisms, underscoring the limitations of current approaches^14^.

One aspect that nonetheless distinguishes GOF and DN mutations is their tendency to spatially cluster within protein structures, in contrast to the more dispersed distribution typically seen for LOF mutations. This phenomenon has been well documented in cancer genetics, where GOF mutations in oncogenes cluster more tightly than the predominantly LOF mutations observed in tumor suppressor genes^21–24^. Similar clustering patterns are now recognized across the broader proteome: GOF-associated variants preferentially localize within defined regions in both sequence and structural space^25–29^. Notably, we recently showed that DN mutations are also characterized by spatial clustering. Using a knowledge-based metric—the Extent of Disease Clustering (EDC)—we demonstrated the ability to identify both GOF and DN disease genes based on variant distributions in PDB structures at proteome scale^14^.

Several computational frameworks have been developed to detect and utilize missense variant clustering at the gene level, particularly in cancer-related genes^24,30–35^. Although both sequence-based and structure-based clustering methods have shown promise, only a fraction of spatial clusters can be captured through sequence analysis alone^36^. A previous VEP tool called DeMAG has attempted to incorporate 3D context into variant-level evaluation by analyzing the spatial neighborhood of known variants, but the targets with available prediction scores are limited^37^. We recently applied a similar concept, in the form of a knowledge-based Euclidean distance metric termed SPDV (Spatial Distance to Disease Variants), to variant-level evaluation of RyR1 mutations^38^. The study demonstrated that SPDV shows comparable performance to dedicated VEP methodologies, but also provides an orthogonal strategy to variant evaluation as it showed different performance levels on variants characterized by distinct RyR1 disease phenotypes.

The advent of high-accuracy structure prediction models, such as AlphaFold2 (AF2), in conjunction with comprehensive variant databases (e.g., ClinVar, HGMD, gnomAD), offers a transformative opportunity to scale structural analyses across the proteome^39–42^. Yet, to date, no VEPs fully capitalize on these resources, and persistent performance biases across variant mechanisms suggest an underutilization of structure-based features that may be particularly informative for non-LOF pathogenicity. In this study, we explore the integration of computational protein structure models and large disease variant datasets to derive heuristic, structure-informed metrics for variant interpretation. We demonstrate that EDC can be systematically applied to AF2-derived models to identify candidate GOF and DN disease genes at scale. Furthermore, we utilize the current knowledge of known pathogenic variant sites to derive the SPDV metric at scale for thousands of disease genes and compare its performance with existing dedicated VEPs. In the general case, SPDV can identify disease variants as effectively as at least half of the tested specialized VEPs, even when only a limited number of disease variant positions are known. However, SPDV performance dramatically improves in high EDC genes, where disease variants spatially cluster, even without either of the metrics utilizing information on benign variant positions. We show that out of the three molecular disease mechanism groups, SPDV most effectively identifies GOF disease variants. Importantly, although the knowledge-based nature of the metric limits its widespread utility as a VEP, its use as an analytical metric enables the identification of a subset of putative GOF disease genes where current VEP methods underperform relative to SPDV, indicating they may be improved by incorporating spatial distance information.

## Results

### Disease variant clustering in high-confidence AF2 model regions allows to identify non-LOF disease genes

As a foundation for our analysis, we identified 3,220 proteins with at least two unique pathogenic missense variant positions, the minimum required to calculate and evaluate clustering or distance-based metrics. Pathogenic variant annotations were sourced from ClinVar^41^ and HGMD^42^. In contrast to our previous work, which primarily relied on experimentally determined structures, we utilized AF2-predicted models^39^ to maximize structural coverage across the proteome, including disordered regions that are typically absent in crystallographic data. To ensure completeness and consistency, we limited our analyses to proteins fully represented within a single AF2 model. Spatial relationships between residues were quantified as Euclidean distances between residue centers of mass, enabling comparison between known disease variant sites and the rest of the protein structure. Per-gene inheritance patterns and molecular disease mechanism annotations were obtained from our prior studies^14,43^.

Previously, we introduced a knowledge-based per-gene clustering metric – Extent of Disease Clustering (EDC) – which effectively distinguishes LOF from non-LOF disease genes when derived using PDB structures^14^. EDC is defined as the ratio between the average shortest distance from non-disease positions to disease positions and the average shortest distance among disease positions themselves (**Figure 1, a**). Higher EDC values reflect increased clustering of pathogenic residues and are enriched among genes associated with non-LOF mechanisms.

**Figure 1.**
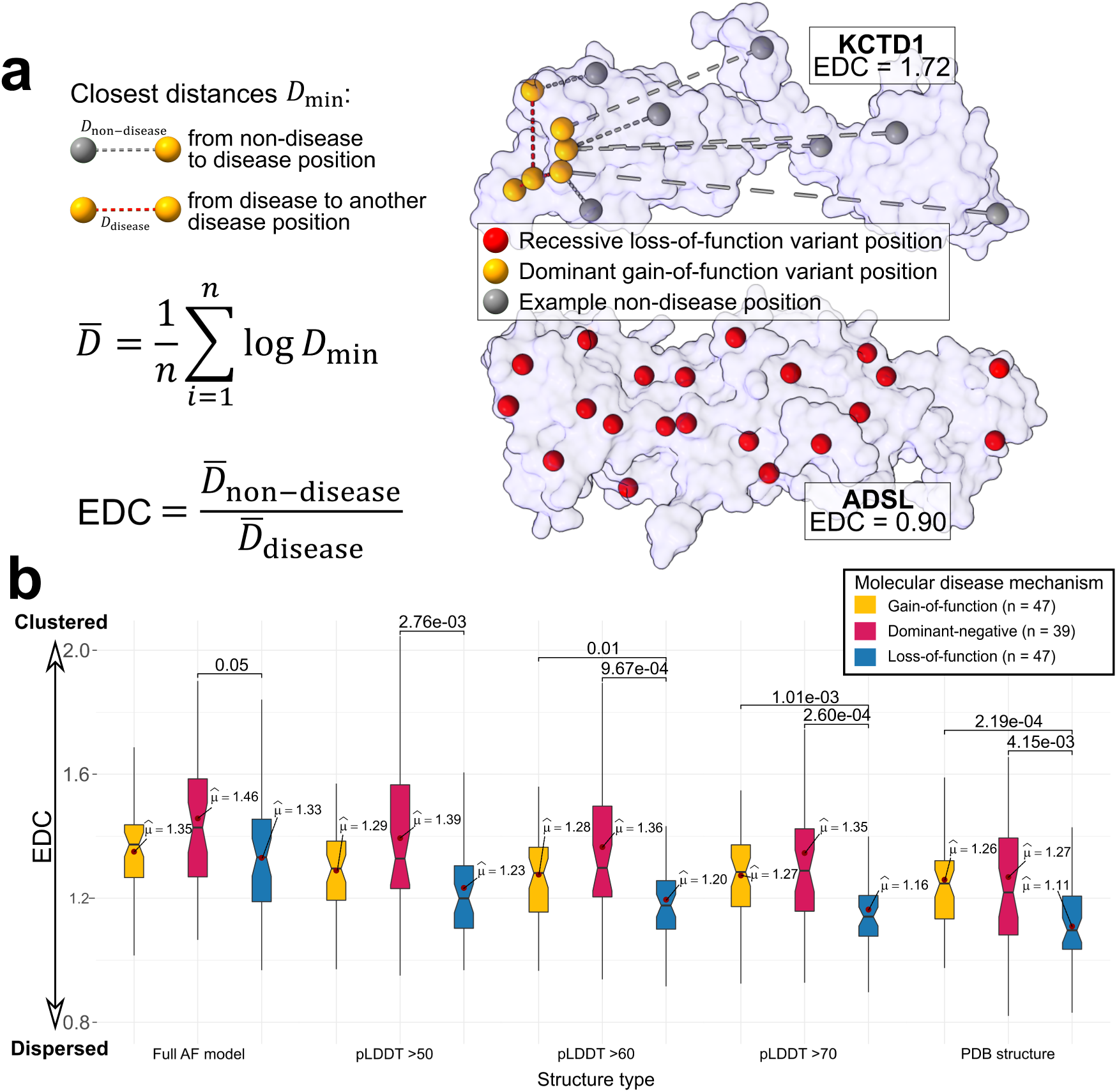
Variant spatial clustering differences between distinct molecular disease mechanisms are most apparent when the Extent of Disease Clustering (EDC) metric is derived using structured protein domains. a – derivation of the EDC metric. EDC is the ratio between two sets of average center-of-mass residue distances within a given protein structure, which both take into account the location of known clinical disease variant positions as a component. The numerator is the average of pairwise distances between disease-harbouring residue positions and all positions without known disease variants. The denominator is the average pairwise distance between all disease positions themselves. An increase in EDC corresponds with an increase in disease variant spatial clustering within a structure. EDC is not influenced by locations of known putatively benign clinical variants. b – Comparison of EDC values derived from PDB structures and AF models filtered to distinct pLDDT thresholds. Boxes denote data within 25th and 75th percentiles, and contain median (middle line) and mean (red dot) value notations. Whiskers extend from the box to furthest values within 1.5x the inter-quartile range. Significance values are derived using two-sided Holm-corrected Dunn’s tests. Sample sizes in the inset indicate the number of genes per group.

However, AF2 models differ from experimental structures in a critical way: they include intrinsically disordered regions, which are generally not resolved in crystallographic PDB entries. The latter predominantly consist of structured domains or disordered segments that adopt ordered conformations upon complex formation. To assess how these low-confidence regions affect EDC performance, we examined the impact of filtering AF2 models by pLDDT, a per-residue confidence metric that inversely correlates with intrinsic disorder^44^.

We recalculated EDC values using three model types: (1) available PDB structures, (2) their corresponding full-length AF2 models, and (3) AF2 models filtered at increasing pLDDT thresholds (>50, >60, and >70). As shown in **Figure 1, b**, EDC values derived from full AF2 models failed to significantly differentiate GOF from LOF genes and showed only marginal separation between DN and LOF genes. However, as low-confidence residues were incrementally excluded, the EDC values for LOF genes declined more sharply than those for GOF or DN genes, revealing more distinct clustering differences. Notably, filtering at pLDDT >70 produced EDC values closely resembling those derived from PDB structures, while retaining broader structural coverage. Such a drastic difference most likely results from the pruning of disordered ‘fuzzball’ models, which would lead to perceived artificial clustering of LOF variants. However, upon removing low-confidence residues, the clustering still remains elevated in GOF and DN disease genes.

Based on these observations, we adopted a pLDDT >70 threshold for all subsequent EDC analyses. Although this filtering reduced the number of proteins with sufficient mapped disease variants, we retained a robust dataset of 2,830 genes, each with at least two pathogenic variants in high-confidence regions. Applying an updated set of molecular disease mechanism annotations^45^, we confirmed that EDC remains effective in distinguishing LOF from non-LOF genes at scale. This distinction was particularly pronounced among autosomal dominant (AD) disease genes (**Supplementary** Figure 1), which are enriched for GOF and DN mechanisms.

### A simple spatial distance metric can identify pathogenic missense variants as effectively as half of the tested VEPs

Building on the gene-level utility of the EDC metric, we sought to evaluate whether spatial knowledge of disease-associated positions could also enhance variant-level pathogenicity prediction. In prior work on RyR1, we introduced a knowledge-based distance metric that showed mechanism-specific performance trends^38^. However, the performance of this metric has not yet been explored at scale.

Spatial Proximity to Disease Variants (SPDV) represents the Euclidean distance between the center of mass of a residue of interest and the residue at the nearest known pathogenic variant location within the same protein structure (**Figure 2**). For residues that already harbor a pathogenic variant, the distance to the next-nearest disease position is used, ensuring an unbiased annotation, as disease positions are not informed of their own classification. To increase robustness to outliers and isolated disease positions, we also computed SPDV as an average across multiple nearest pathogenic positions. Under this framework, lower SPDV values indicate closer proximity to known disease positions, and thus a potentially higher likelihood of pathogenicity.

**Figure 2.**
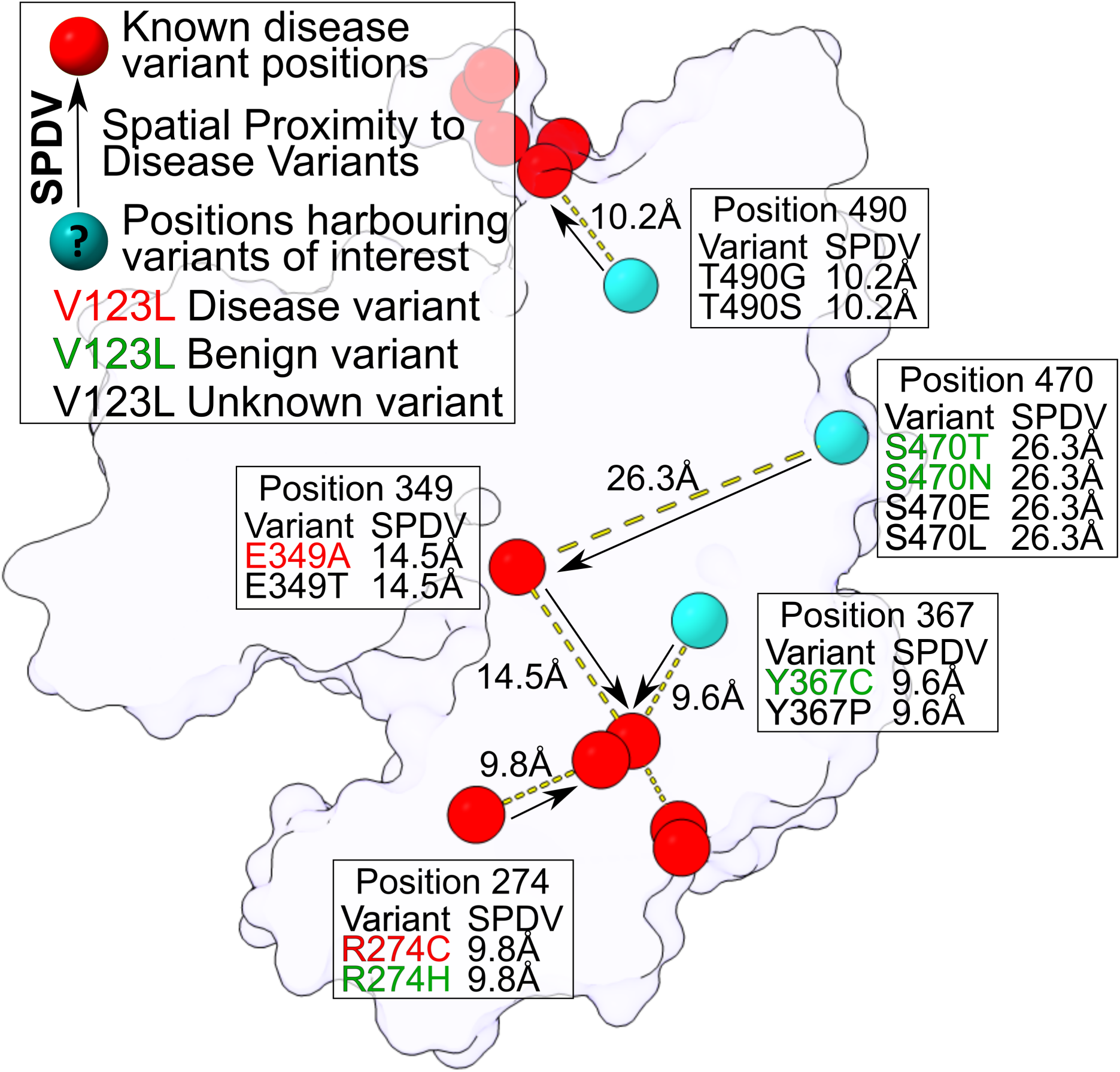
Derivation of the Spatial Proximity to known Disease Variants (SPDV) metric using AlphaFold protein structures. Analogously to EDC, SPDV is computed solely from disease variants data, without incorporating information about the positions of clinically benign variants. SPDV represents the Euclidean distance from a residue position of interest to the closest position harbouring disease variants. Distances are calculated between wild-type residue centers of mass. As a position-level metric, all variants at a given position are annotated with the same SPDV value, regardless of underlying classification. Self-positions are excluded from distance calculations, and as such, positions that themselves harbour disease variants are not influenced by their own classification. The figure example depicts SPDV_1, a distance to a single closest disease position for each residue. SPDVs can also be derived using an average of distances to a number of closest disease positions (SPDV_5 is an average of distances to five closest unique positions, etc.).

To calculate and evaluate SPDV at scale, we compiled a dataset of pathogenic missense variants from ClinVar and HGMD, as well as putatively benign ClinVar and gnomAD^40^ mutations. As gnomAD contains some heterozygous or incompletely penetrant variants that may be damaging, we excluded any pathogenic variants present in ClinVar and HGMD from our putatively benign variant dataset. We included only protein targets with at least two distinct pathogenic variant positions and structures fully covered by a single AF2 model, resulting in 3,073 proteins annotated with SPDV scores derived from up to 40 nearest pathogenic positions (with SPDV_1 indicating the distance to a single closest disease variant position and SPDV_40 indicating a value resulting from the averaging of distances to 40 nearest positions).

To optimize our use of SPDV, we first aimed to explore how inclusion of disordered regions and averaging of distances to multiple disease positions may influence pathogenicity prediction performance. SPDV was assessed via per-gene receiver operating characteristic (ROC) analysis, with area under the curve (AUC) as the primary metric. As shown in **Supplementary** Figure 2, filtering out disordered residues (using pLDDT) decreased SPDV performance—attributable to the fact that benign variants are frequently located in peripheral, disordered regions. As such, removing disordered residues reduces the perceived performance, as benign variants that are easy to identify due to their peripheral location and thus higher SPDV values are removed. Accordingly, we used SPDV values derived from full-length AF2 models, which ensure maximal variant coverage and are most comparable to sequence-based VEPs that likewise include disordered regions.

Averaging SPDV across multiple disease positions further improved predictive performance. **Figure 3, a** demonstrates that using a single closest distance (SPDV_1) already achieves an AUC of ∼0.69, with subsequent averages of up to 25 peaking at 0.75 AUC. However, including more neighbors reduced gene coverage due to limited variant data. Based on this trade-off, we selected SPDV averages across up to 5 positions for further analysis.

**Figure 3.**
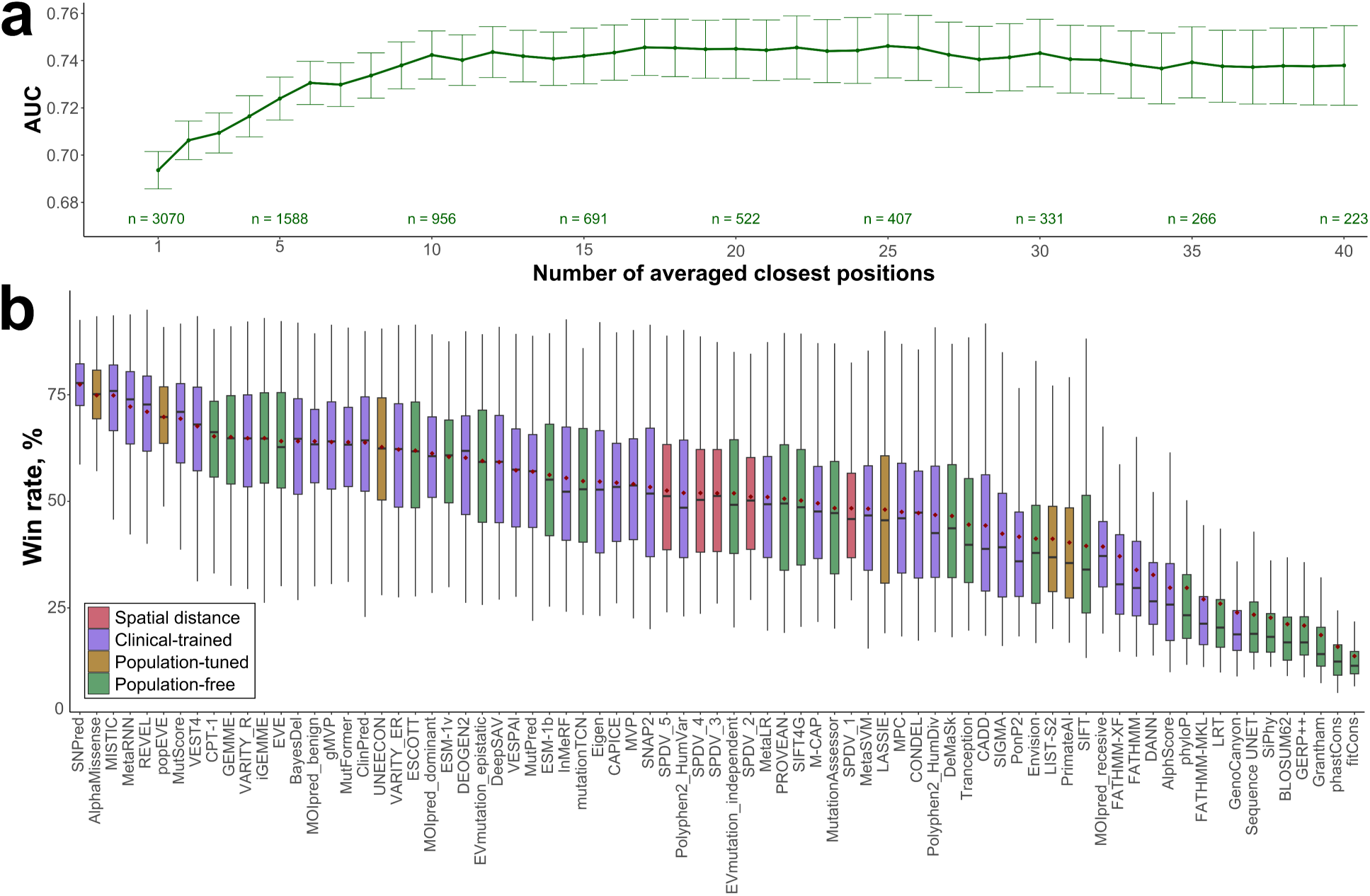
SPDV outperforms half of the tested dedicated variant effect predictors on autosomal dominant disease genes, even with knowledge of only a few disease positions. **a** – Median per-gene performance and sample size comparison of SPDV values derived through averaging up to 40 distances to closest disease positions. Sample sizes indicate the number of genes per group for the given number of averaged positions. Bars represent the 95% confidence interval for the median, and extend 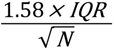 (McGill et al., 1978). **b** – Performance comparison of SPDV metrics and dedicated VEPs on 849 autosomal dominant disease genes. Win rates were calculated using a pairwise AUC comparison scheme (see Methods). Boxes denote data within 25th and 75th percentiles, and contain median (middle line) and mean (red dot) value notations. Whiskers extend from the box to furthest values within 1.5x the inter-quartile range.

Due to a higher protein coverage and an abundance of the putatively benign gnomAD variants we also speculated that a reverse metric, the distance to the closest benign disease variant position, could serve as a useful alternative predictor. However, performance was inconsistent. As shown in **Supplementary** Figure 3**, a**, averaging over one or two benign variants yielded near-random AUCs (∼0.5), while greater averaging resulted in inverse behavior, with AUCs declining to ∼0.33. This counterintuitive result likely reflects the distinct spatial distributions of benign and pathogenic variants. Specifically, ∼70% of benign variants occur on protein surfaces, compared to only ∼41% of disease-associated ones (**Supplementary** Figure 3**, b**). As a result, pathogenic variants often reside closer to peripheral benign variants than benign variants do to each other across the surface, leading to inflated average distances and reduced discriminative power. Although the inverse benign metric peaked at an AUC of ∼0.67, it still underperformed compared to SPDV, and its signal appears largely driven by interior vs. surface variant localization.

To more rigorously benchmark SPDV against dedicated VEPs, we focused on a subset of 849 AD disease genes, which represent higher confidence disease targets and are enriched in diverse molecular disease mechanisms. We compared SPDV against 72 methodologically diverse VEPs, all of which have been recently benchmarked against deep mutational scanning and clinical variants datasets, using pairwise win-rate comparisons based on AUC (see Methods)^46^. These predictors span a range of methodological categories, including: (1) supervised predictors directly trained on clinical variants (clinical-trained), whose performance is likely overestimated due to being exposed to at least some of our benchmarking set variants; unsupervised methodologies, such as language models or conservation-based predictors, some of which were (2) fine-tuned on population frequency data (population-tuned), or (3) are mostly unbiased (population-free) and the most directly comparable to SPDV.

As **Figure 3, b** illustrates, SPDV metrics with distance averages of 2–5 nearest neighbors performed competitively, winning ∼51–53% of pairwise comparisons. This performance is on par with or exceeds several widely used tools, including PolyPhen-2, the epistasis-independent EVmutation model, and SIFT4G, and trails only slightly behind state-of-the-art models such as ESM-1b. Even the simplest SPDV_1 metric achieved a 48% win rate average, while being subjected to the largest number of pairwise comparisons against other VEPs.

Notably, SPDVs’ competitive performance is particularly striking given its simplicity: it is a position-based metric, assigning the same score to all variants at a given residue, regardless of the specific substitution. Despite this limitation, SPDVs remain comparable to many sophisticated VEPs. Interestingly, SPDV also exhibited greater heterogeneity in win rates across genes, suggesting that it may be more effective for specific variant classes or mechanisms.

### SPDV is highly effective at identifying GOF disease variants

We have previously shown that variant spatial clustering in 3D structural space shows distinct tendencies among dominant disease genes characterized by different molecular mechanisms, such as gain-of-function (GOF), the dominant-negative (DN) effect and haploinsufficient loss-of-function (LOF)^14^. In particular, non-LOF variants tend to cluster, whereas LOF variants are more dispersed, consistent with their typical impact on protein destabilization and degradation. Building on this observation, we sought to determine whether spatial clustering could enhance variant-level pathogenicity prediction using SPDV.

We repeated our SPDV benchmark, but this time stratifying the analysis by molecular mechanism using a previously curated set of high-confidence annotations for GOF, DN, and haploinsufficient LOF disease genes^43^. To minimize potential bias, we restricted comparisons to 27 unsupervised, population-free VEPs, which are less prone to data circularity. **Figure 4** shows that SPDV_2 emerges as the third-best performing predictor for GOF genes, achieving a mean win rate of 67.5%, only 0.69% behind the top-ranking method, iGEMME. Even SPDV_1, despite being the simplest version of the metric, outperformed most other VEPs – achieving a 62.5% win rate, surpassing tools such as ESM-1b and EVmutation.

**Figure 4.**
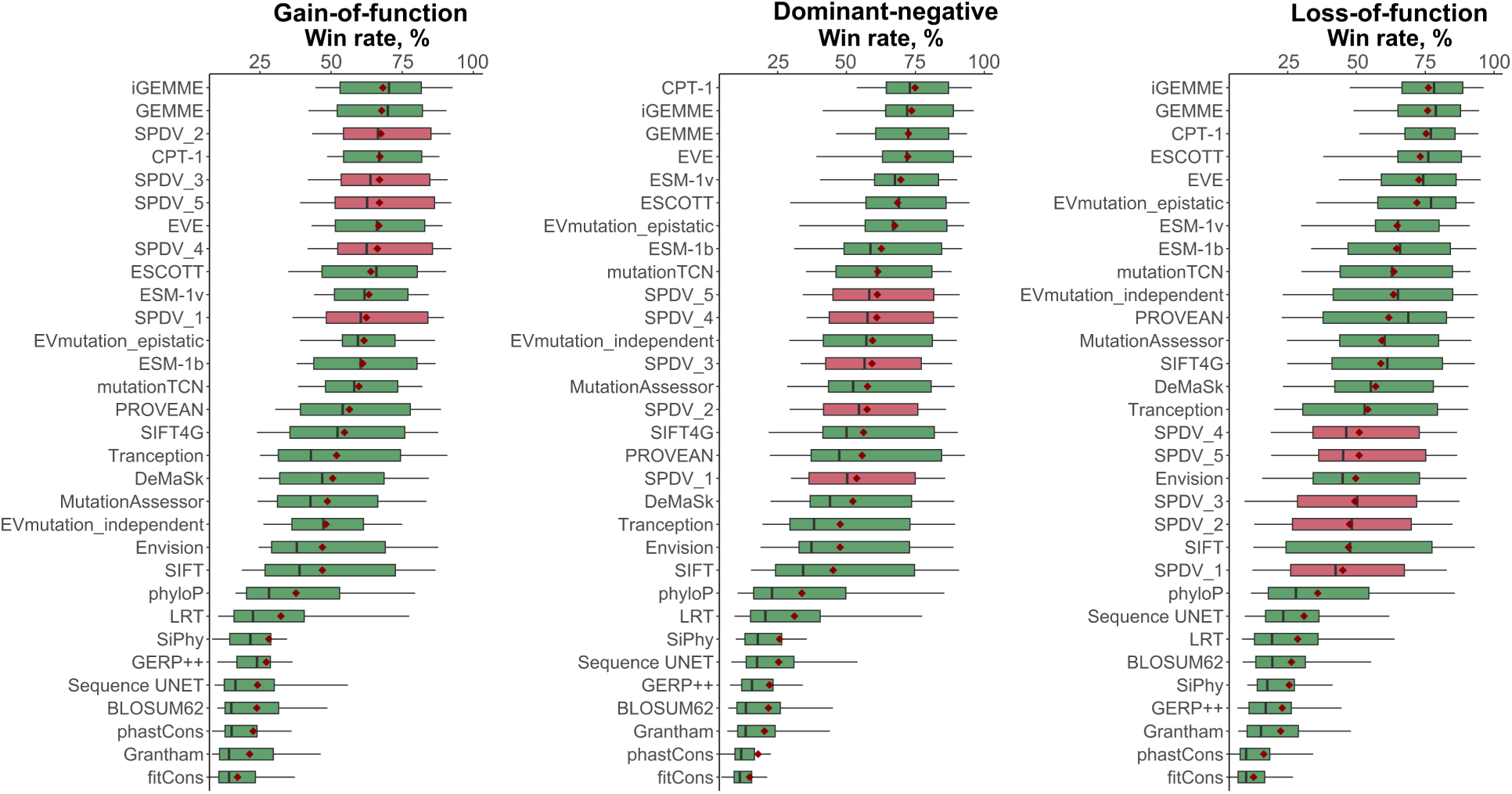
SPDV particularly excels at identifying GOF disease variants. Win rates were calculated using a pairwise AUC comparison scheme (see Methods). Boxes denote data within 25th and 75th percentiles, and contain median (middle line) and mean (red dot) value notations. Whiskers extend from the box to furthest values within 1.5x the inter-quartile range.

SPDV metrics show worse performance for DN and then for LOF genes, with the best win rates at ∼61.2% and 51%, respectively, indicating that spatial variant clustering is the most leverageable for GOF disease genes. While it has been previously shown that most existing predictors have a negative performance bias against GOF and DN variants^14^, the simple heuristic SPDV metric favours non-LOF disease genes specifically.

Our previous mechanism-centric exploration of VEP performance identified that disease variants in genes associated with DN and GOF mechanisms were universally underpredicted by VEPs of the time. However, in recent years an abundance of new population-free VEPs have been released, which may have brought potential improvements in this regard. Indeed, **Supplementary** Figure 4 demonstrates that the top methods we have explored in this work now show an almost equivalent performance level between DN and LOF disease variants. Interestingly, despite their favourable general performance, the newest VEPs still underpredict GOF variants in comparison to LOF disease, indicating an alternative approach may be needed to close the performance gap.

Although gene-level mechanism annotations are unavailable for most proteins, we have identified EDC as a useful quantitative proxy for identifying putative non-LOF disease genes. Given the strong mechanistic link between variant clustering and SPDV performance (**Figure 4**), we evaluated the applicability of SPDV through the scope of protein-specific EDC values across up to 2,830 genes. As shown in **Supplementary** Figure 5, there is a strong positive correlation between the performance of SPDVs and EDC, with the highest correlation being found for SPDV_1 (Spearman’s rho 0.83), suggesting that EDC may be predictive of genes where SPDV will be the most effective.

Using this insight, we sought to identify whether there is an EDC threshold at which SPDVs consistently exceed all other VEPs for a given subset of genes. Using only genes with at least 5 known distinct disease positions, we iteratively filtered the dataset based on increasing EDC thresholds and calculated pairwise predictor win rates. We found that while iGEMME and CPT-1 dominated the pairwise comparisons initially, SPDV_2 becomes the top-ranking predictor at an EDC of ∼1.355, consistently beating other SPDVs and all tested population-free VEPS throughout the rest of the range, starting with a mean win rate of 74.8% (**Figure 5**, **Supplementary** Figure 6).

**Figure 5.**
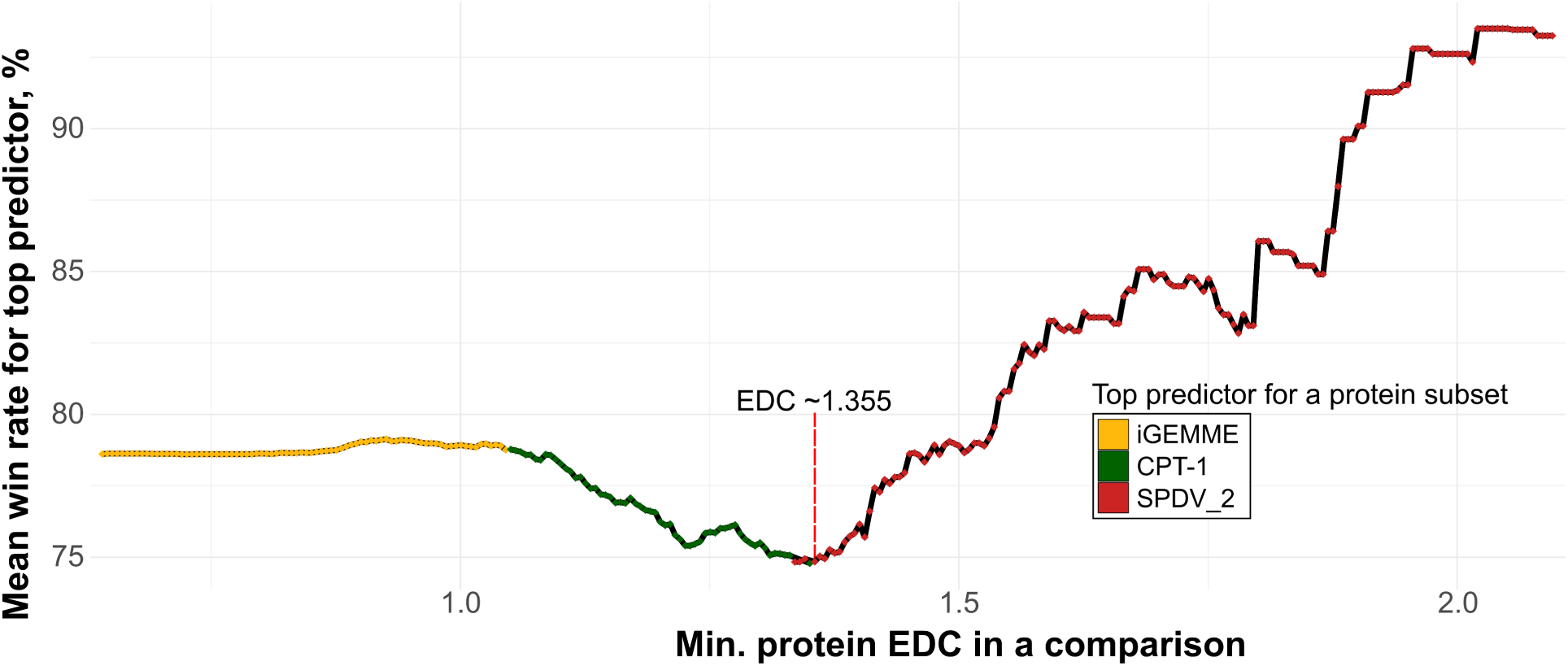
SPDV_2 consistently overtakes other VEPs in disease variant idenfitication performance, after a low-point of single-VEP dominance and increasing extent of overall disease position clustering. Win rates were calculated using a pairwise AUC comparison scheme (see Methods). The benchmark was carried out starting from a set of 1,875 disease genes with at least 5 distinct disease positions, annotated with EDC values. An EDC step of 0.05 was used to iteratively filter out genes to assess the performance of the five SPDV metrics and other population-free VEPs in a win rate competition on each protein subset. The red line marks the point of EDC ∼1.355 where SPDV_2 consistently (for all subseqeuent thresholds) becomes the best overall predictor.

While iGEMME appears to consistently outperform other predictors in the low EDC range that includes most LOF-associated disease genes, it eventually gives way to CPT-1, which appears to become the top predictor while also demonstrating a reduced overall win rate. This observation indicates that with increasing EDC, the ability of the top predictor to perform consistently well across all proteins diminishes, with some VEPs appearing to outperform it on select proteins. This trend reverses at the threshold of EDC ∼1.355, where increasing variant clustering also drives SPDV performance. The noted threshold encompasses 371 high-EDC genes, 272 of which were not currently annotated as GOF or DN disease targets in our data, suggesting they may represent previously unrecognized non-LOF disease genes.

While neither EDC nor SPDV metrics utilize any information on benign variant positions, we recognize a spatial metric may benefit from the tendency of peripheral disordered regions to contain more neutral variants, which SPDV would have an advantage of identifying due to long Euclidian distances, compared to conventional VEPs. A recent work has also highlighted that current VEPs perform inconsistently between ordered and disordered regions, with noted underperformance in the latter ^47^. To control for this, we repeated the EDC step analysis, but this time only considering residue positions above 70 pLDDT, the per-residue AlphaFold modelling confidence metric which is highly correlated with structural order^48^. Interestingly, we show that SPDVs still consistently exceed other VEPs in performance after a certain threshold, although at a slightly higher EDC of ∼1.369 when only considering ordered positions (**Supplementary** Figure 7). In this instance, SPDV_2 also was not the top SPDV consistently, with SPDV_1 temporarily overtaking it in the ∼1.52-1.55 EDC range.

To more quantitatively evaluate the impact of variants in disordered regions we contrasted the win rates of all SPDVs and population-free VEPs when derived on full models *vs* just the variants occurring in structurally ordered (70 > pLDDT) positions, and derived Δ win rates, with negative values indicating a higher win rate on full model datasets, while positive values showed the metrics had better performance when only the ordered positions were considered (**Supplementary** Figure 8). We considered two protein subsets, with the first containing 1724 genes with at least 5 distinct disease positions, while the second, in addition, was filtered down to only those with a disease variant clustering extent above 1.355, identified previously. Interestingly, we find that while overall SPDVs are the most advantaged by the presence of variants in disordered regions, the high-EDC protein subset is not affected to any considerable degree, demonstrating mean Δ win rate values for SPDVs at around 0. Thus, the exceptional SPDV performance in the latter niche, exceeding all other population-free methods, is driven purely by the phenomenon of disease variant clustering itself, and SPDV serves as a diagnostic metric to identify targets where current VEP methodologies demonstrate blind spots. These findings highlight the utility of spatial disease variant clustering and the potential of combining EDC and SPDV as a framework to both identify candidate pathogenic variants and infer underlying disease mechanisms in uncharacterized genes.

## Discussion

Despite the exceptional general performance of modern variant effect predictors (VEPs), they still demonstrate inconsistent performance between disease variants arising through distinct molecular mechanisms, tending to perform better on loss-of-function (LOF) disease genes. In this study we have further built upon the concept of the knowledge-based SPDV^38^ (Spatial Proximity to Disease Variants) metric by implementing it at scale for thousands of protein targets. We demonstrated its unique utility in accurately capturing the pathogenicity of non-loss-of-function (non-LOF) disease variants, but also as a diagnostic metric that allows to identify protein subsets where current VEPs underperform compared to a simple heuristic metric.

In our benchmark, using the distance to just a single closest known disease variant position within a protein structure surpasses at least a third of the 72 dedicated variant effect predictors (VEPs) in dominant disease genes, while SPDVs derived through averaging of distances to 2-5 positions exceeded the performance of half of the methodologies. However, the utility of SPDV becomes especially evident when focusing on gain-of-function (GOF) disease genes. In this context, SPDV performance essentially matches that of the top-ranking population-free VEPs, highlighting its potential as a simple, interpretable, and mechanism-aware feature to close the predictive performance gap between LOF and non-LOF disease variants.

While SPDV directly utilizes data on clinically relevant disease variant locations, each distance measurement is an unbiased empirical observation and unaffected by the underlying classification for the position of interest. Furthermore, the disease variant identification performance of SPDV, as measured by AUC or the win rate, in part depends on the spatial distribution of clinically benign variants. However, neither SPDV, nor EDC, are trained or derived with any knowledge of benign clinical variant positions, leading us to focus only on fair comparisons against population-free VEPs for most of our analyses. Population-free VEPs are unsupervised methods that have not been fine-tuned using variant frequencies observed in populations and are less affected by data circularity concerns, compared to clinical-trained or population-tuned VEPs which may have been trained or tuned on our benchmarking data^46,49^.

Further, using the Extent of Disease Clustering (EDC) metric to prioritize target proteins exceeding a certain degree of variant clustering, we demonstrated SPDV_2 becomes the overall best predictor against the tested unbiased population-free VEPs. While top VEPs are consistently starting to utilize structural information in prediction pipelines^1^, and spatial clustering has been previously leveraged for disease gene and hotspot identification^24,30–34,37^, no existing method has implemented spatial proximity to disease variants at scale to directly improve variant-level classification across molecular disease mechanisms^14^. While target prioritization using EDC might appear circular due to SPDV essentially being a position-level derivative of EDC, neither of the metrics is derived with any knowledge of benign variant locations. As such, a target with a high EDC value may display lower than expected SPDV performance due to benign variants also occurring at or proximal to disease variant positions. However, despite this possibility, SPDV performance demonstrates a high correlation with EDC, indicating benign variants tend to not occur at spatial disease variant clusters. Our results demonstrate that both EDC and SPDV represent powerful, complementary metrics that can enhance future predictive frameworks, or serve as diagnostic metrics to probe protein niches where current VEPs could be further improved.

To expand applicability of SPDV to proteins lacking sufficient disease variant data, we investigated an alternative approach: measuring the distance to the nearest benign variant, leveraging the abundance of population-level variation captured in resources like gnomAD. While this ‘inverse’ SPDV metric offers broader theoretical coverage, its predictive performance was limited, with a maximum AUC of only ∼0.67 when averaging distances to the 40 closest benign positions. We suspect this modest performance stems mostly from the bias in disease variant datasets, which are enriched for LOF disease proteins and mutations that typically localize to structured protein cores. In contrast, benign variants more often occur on the protein surface. As a result, when averaging distances to many benign sites, pathogenic variants located in the interior may appear, on average, closer to surrounding surface-exposed benign variants than benign variants are to each other across the protein surface, leading to predictive performance relying solely on spatial distribution.

We also demonstrated that using full AlphaFold2 (AF2) models, which include disordered regions marked by low pLDDT scores, can significantly influence the perceived performance of distance-based pathogenicity prediction metrics. These disordered regions, often peripheral to the well-structured protein core, tend to harbor a higher density of benign variation. Consequently, variants in these regions become easier to distinguish based on spatial separation alone. This effect does not detract from the utility of SPDV, particularly as our benchmark comparisons were conducted against sequence-based VEPs that use full-length proteins. Rather, it underscores the heuristic nature of the metric, which intentionally leverages the differential spatial distributions of variant classes. However, it also highlights key distinctions between AF2 models and experimental PDB structures, especially for structure-based applications, where future work may benefit from selectively including or excluding disordered regions depending on the predictive task (e.g., structure-based stability assessments may suffer with disorder included).

While we show that structural distance to known disease positions can effectively improve the prediction of non-LOF variant pathogenicity, we acknowledge that this feature is inherently auxiliary. It is most useful in genes with a sufficient number of annotated pathogenic variants and is particularly suited to improving identification of GOF and DN mechanism variants and genes. Beyond the requirement for known disease positions, other challenges may include cases where the target contains mixed-mechanism phenotypes, for instance across separate domains, or pathogenic variant distribution patterns that are more visible in the context of a multi-subunit complex assembly. Our analyses were limited to proteins for which complete structural models were available in a single AF2 file, excluding some larger and fragmented targets. Future work should explore the use of spatial distance at the domain level and across entire protein assemblies, ensuring non-LOF disease variant clusters are identifiable throughout the noise of spatially distributed LOF variation.

Ultimately, SPDV and EDC represent promising steps toward correcting the LOF-centric bias that persists in most current variant interpretation frameworks, with the spatial disease variant distribution patterns being just one possible avenue to distinguish the distinct mechanisms. We hope this work encourages the development of additional simple, interpretable, mechanism-aware metrics that can enhance variant effect prediction—particularly in under-annotated or under-recognized GOF and DN disease genes. The SPDV scores we provide for all possible missense substitutions across eligible proteins may serve as a useful resource for variant reclassification efforts and mechanistic exploration in both clinical and research settings.

## Methods

### Derivation of the structural variant dataset and annotations

Variant and gene clinical classification, mechanism and inheritance annotations, derived distance metrics, variant effect predictions and spatial clustering values are accessible at https://doi.org/10.17605/OSF.IO/6PTFK.

The pathogenic and benign mutation lists were compiled using ClinVar (downloaded after 2024.05.01), HGMD and gnomAD (v4.1.0) databases. ‘Pathogenic’ and ‘likely pathogenic’ ClinVar variants, as well as an HGMD-based dataset of disease variants from a recent work by Bayrak *et al.*^13^ were used as our pathogenic variant and position dataset. The putatively benign mutation set was assembled from ClinVar ‘benign’, ‘likely benign’ and gnomAD mutations. gnomAD mutations were not filtered based on allele frequency, as the vast majority of the gnomAD database is comprised of rare or singleton variants. However, we tried to address the existence of incompletely penetrant or heterozygous disease variants in the dataset by filtering out any variants from the disease set, and by deriving a higher confidence subset consisting only of OMIM autosomal dominant disease genes.

To increase the variant structural coverage as the primary source of protein structures we chose AlphaFold2^39^ models, mapping the variants based on UniProt^50^ sequence position annotation. The AlphaFold2 model dataset was downloaded on 2021.07.27 from https://alphafold.ebi.ac.uk/. For the initial dataset, targets with at least 10 individual known disease variant positions were chosen, and we only focused on proteins whose entire structure fit within one model file (targets below 2700 amino acids), due to the nature of the distance metric. For our analysis exploring the applicability of AF *vs* PDB structures when deriving EDC, available experimental subunit structures were downloaded from the Protein Data Bank on 2020.08.17. The first biological assembly for each structure was chosen under the assumption it represents the biologically relevant quaternary structure. Mutations were mapped to protein structures in a manner that was previously described^51^, considering polypeptide chains with >90% sequence identity to a human protein over a region of no less than 50 amino acids, which can include structures of non-human origin. In such cases, mutations were only mapped to structures where the residue of interest, as well as its adjacent neighbours, were the same as the human wild-type sequence. If a residue mapped to multiple PDB files, we selected the structure with best resolution followed by largest biological assembly. Only the first structure was extracted from NMR ensembles. Every variant was mapped only to a single residue in a single PDB structure, and residues missing from PDB structures were not considered in the PDB-based analysis. For PDB files containing multiple occupancies of a single residue, only the first occurring entry was selected.

Molecular disease mechanism annotations, which we had previously derived and described in Gerasimavicius *et al.*^14^, were derived through OMIM^52^ gene entry page curation, disease inheritance pattern group and ClinGen^53^ dosage sensitivity annotation. An updated set of inheritances and molecular disease mechanisms was taken from Badonyi & Marsh^43^.

### Distance metric calculation and variant effect predictions

The various types of SPDV (Spatial Proximity to Disease Variants) distance metrics were derived from protein structures using the ‘bio3d’ R package^54^. Euclidean distances from a residue of interest to all disease residue positions in a structure were calculated using amino acid residue centers of mass as the reference point using ‘dist.xyz’ and ‘atom2mass’ functions. The distances were ranked in ascending order, a single or a number of closest distances were retrieved and averaged for a given SPDV metric. In the case of the ‘inverse’ gnomAD-based metric (SPNV – Spatial Proximity to Neutral Variants), closest distances from residues of interest were calculated to known putatively benign variant positions. In the case where the position of interest itself was associated with a disease or benign variant, depending on whether the closest-disease or closest-benign metric was being calculated, the distance was calculated to the next nearest relevant non-self position, respectively. We provide a Colab notebook to allow easy calculations of SPDVs for all amino acid residue positions in an input structure, given a list of known disease variant positions. The code is available at https://github.com/lgeras/SPDV.

VEP classifications and prediction values were obtained from the recent benchmark by Livesey & Marsh^55^.

### Spatial disease variant clustering

The Extent of Disease Clustering (EDC) metric was first derived in Gerasimavicius *et al.*^14^. The metric is based on the proximity of each protein residue to a known pathogenic variant at another residue in the structure. For each residue, considering only monomeric subunits, we calculated the center of mass distance *D* to all other residues with a known ClinVar disease mutation, and the closest distance *D _min_* was selected. We calculated the average of the log distance (*D̄*) for all disease residues, and all non-disease residues separately (Equation 1).

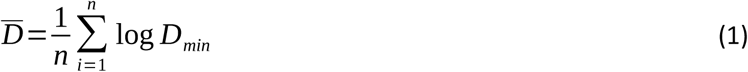

The final clustering metric, which we termed ‘Extent of Disease Clustering’ (EDC), is presented as the ratio of the two values (Equation 2):

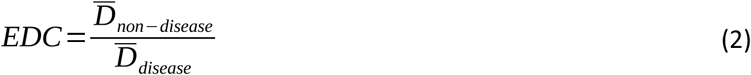

Thus, a value greater than one indicates that the sites of disease mutations tend to be closer to the sites of other disease mutations than non-disease residues tend to be to the sites of disease mutations.

### Statistical testing

ROC analyses were carried out using the ‘pROC’^56^ package, with AUC curve differences being statistically assessed through the DeLong method using ‘roc.test’. In the VEP ROC analysis, case-control direction was implicitly set individually for each predictor. Line plot bars represent the rough 95% confidence interval for median values, as described by McGill *et al.*^57^. Pairwise statistical comparisons between group EDC values were carried out using Dunn’s test implementation in the R ‘ggstatsplot’^58^ package, with the *p*-values for comparisons involving more than two groups being adjusted through Holm’s multiple comparison correction^59,60^.

VEP win rates were calculated based on a scoring system, where for every VEP pair over all given proteins, their AUCs for a common variant subset within that protein were compared, and the winning VEP was assigned 1 point, while a close draw led to both VEPs being assigned 0.5 points. In the end, the win rate for each VEP pair was derived by dividing the scores by the number of successful comparisons that were carried out for a pair over all available genes. VEPs for Figure 3, b were filtered out if they did not have predictions for at least 40% of the genes, compared to the VEP with the most gene coverage. Subsequent win rate calculations involved pre-filtering of VEPs that were not involved in at least one third of the number of pairwise comparisons as the VEP with most comparisons. Mean win rates were derived by averaging over all win rates for a given VEP against each other VEP, individually, across all genes.

The EDC threshold analysis to identify EDC values where SPDVs become top predictors were carried out by taking the minimal EDC value in the dataset and iteratively filtering down the dataset to genes that contain at least that EDC value, with an EDC step value of 0.005. Average win rates were then calculated from the resulting gene data and a ranking was established. Iteration through EDC steps was stopped when the number of viable genes dropped below 10 in each instance. This analysis was performed only on proteins with at least 5 distinct positions harbouring disease variants.

To assess the SPDV performance impact of easily distinguishable neutral variants occurring in disordered peripheral regions of the structures, we repeated the win rate analysis with only disease and benign variants from structured regions (pLDDT>70). This analysis was performed on an updated list of proteins, requiring at least 5 distinct disease positions in ordered regions. The difference in average win rates was derived on a per-gene basis, subtracting the win rate calculated for the full model from the one derived only using variants from structured residues.

## Data & code availability

The data generated in this study have been deposited in the OSF database at https://doi.org/10.17605/OSF.IO/6PTFK. The Google Colab notebook to calculate SPDV values is available at https://github.com/lgeras/SPDV.

## Acknowledgement

This project was supported by funding from the European Research Council (ERC) under the European Union’s Horizon 2020 research and innovation programme (grant agreement No. 101001169) and by funding from the Medical Research Council (MRC) Human Genetics Unit core grant (MC_UU_00035/9).

---

**Supplementary Figure 1.**
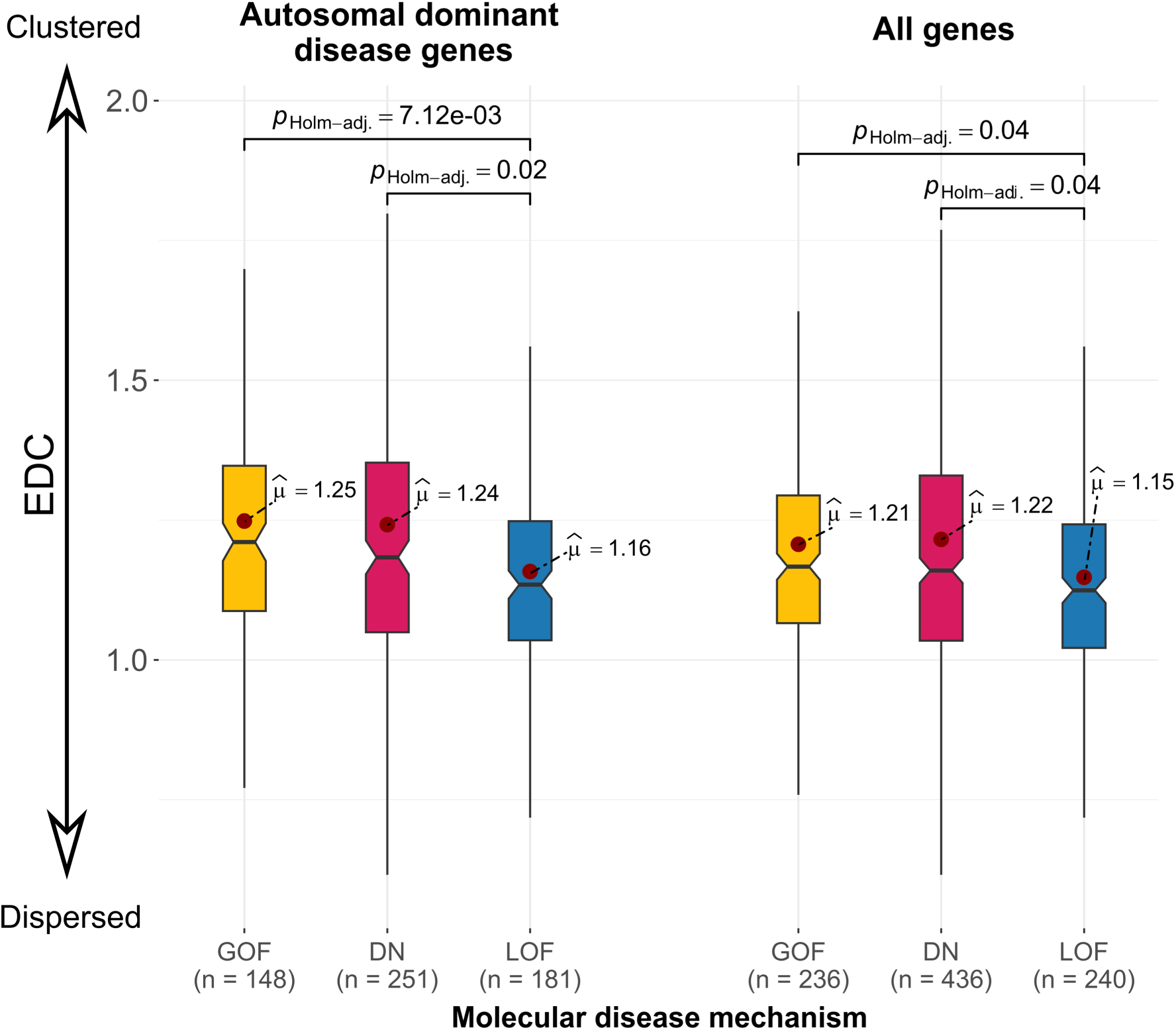
EDC values derived at scale maintain their discriminatory power, especially for autosomal dominant disease genes. Boxes denote data within 25th and 75th percentiles, and contain median (middle line) and mean (red dot) value notations. Whiskers extend from the box to furthest values within 1.5x the inter-quartile range. Sample sizes indicate the number of genes per group.

**Supplementary Figure 2.**
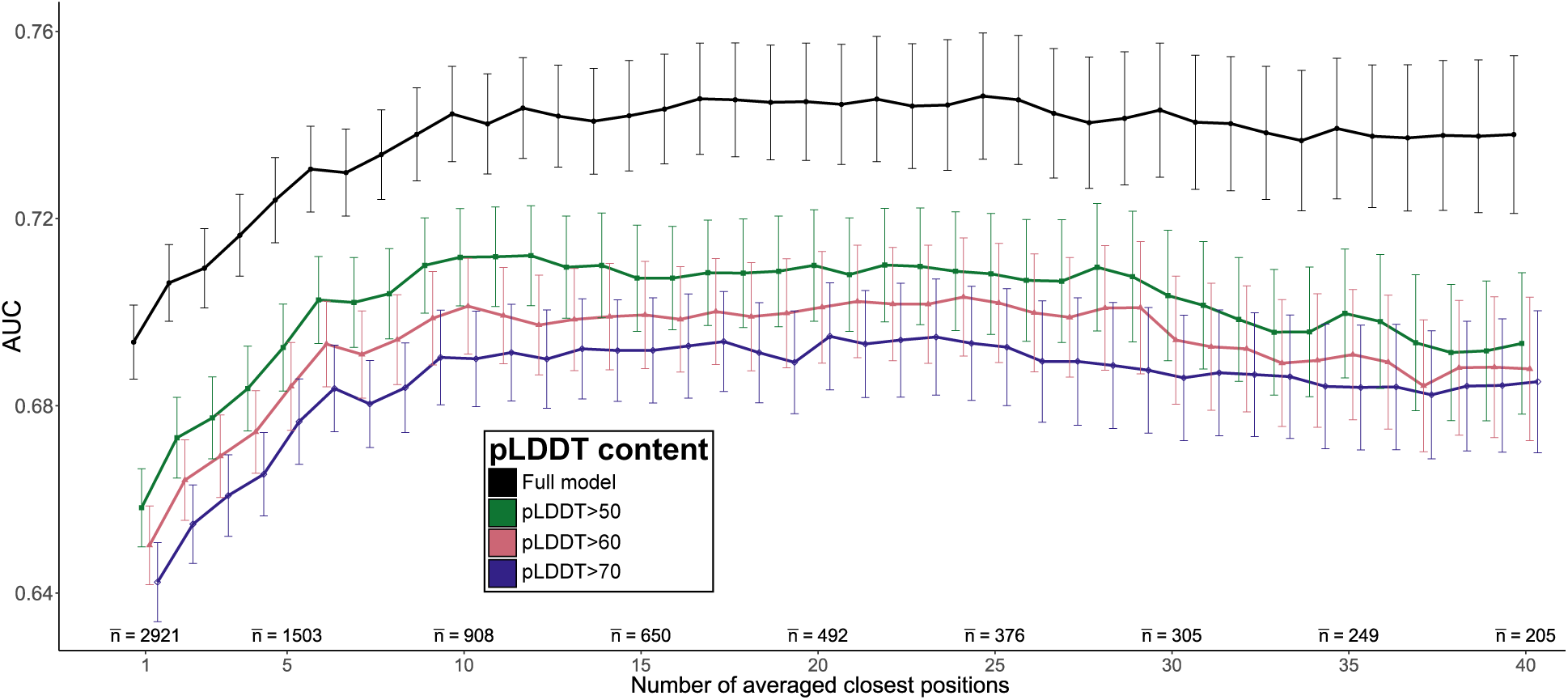
Taking into account variants from highly disordered regions increases the perceived pathogenic variant identification performance of SPDV. Line plot points indicate the median of per-gene area-under-the-curve (AUC) values from receiver operating characteristic analyses comparing between pathogenic and putatively benign ClinVar, HGMD and gnomAD variants. Points for the same position count are staggered per group. Points that show no confidence interval overlap with other groups for a given averaged position count are highlighted with a yellow outline. For these positions, bars represent the 95% confidence interval for the median, and extend 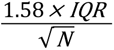 (McGill et al., 1978). Sample sizes indicate the average number of genes per position group.

**Supplementary Figure 3.**
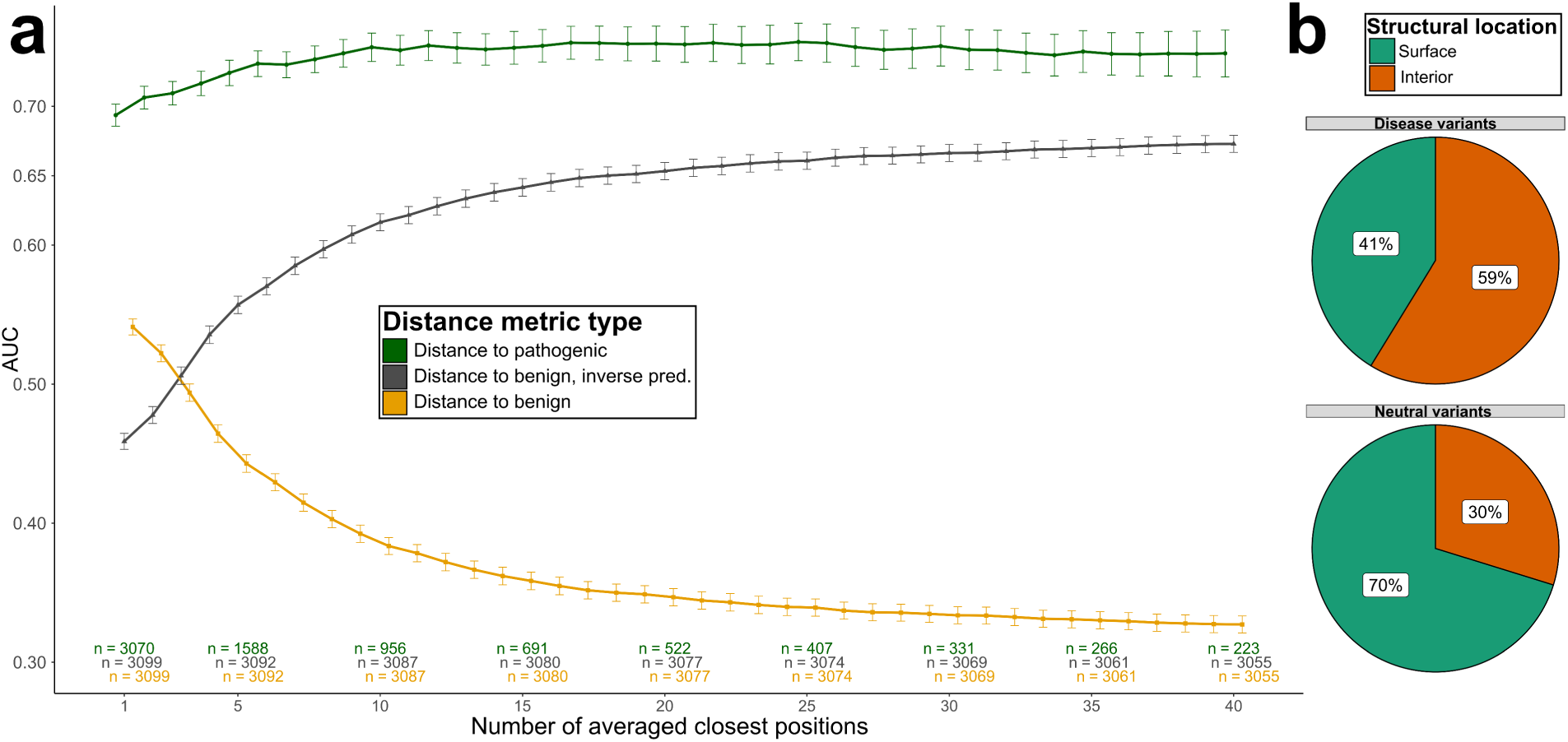
Distance to known disease (SPDV) and not to benign variant positions is most predictive of pathogenicity, and achieves high performance with knowledge of just several closest positions. **a** – Performance of SPDV and an analogous distance to benign variant metric. **b** – Structural location distribution differences between disease and benign variants. Line plot points indicate the median of per-gene area-under-the-curve (AUC) values from receiver operating characteristic analyses comparing between pathogenic and putatively benign ClinVar, HGMD and gnomAD variants. Bars represent the 95% confidence interval for the median, and extend 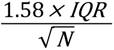 (McGill et al., 1978). The grey datapoints represent inverted AUC values for the metric using closest positions to known benign variants (grey). Sample sizes indicate the number of genes per group for the given number of averaged positions.

**Supplementary Figure 4.**
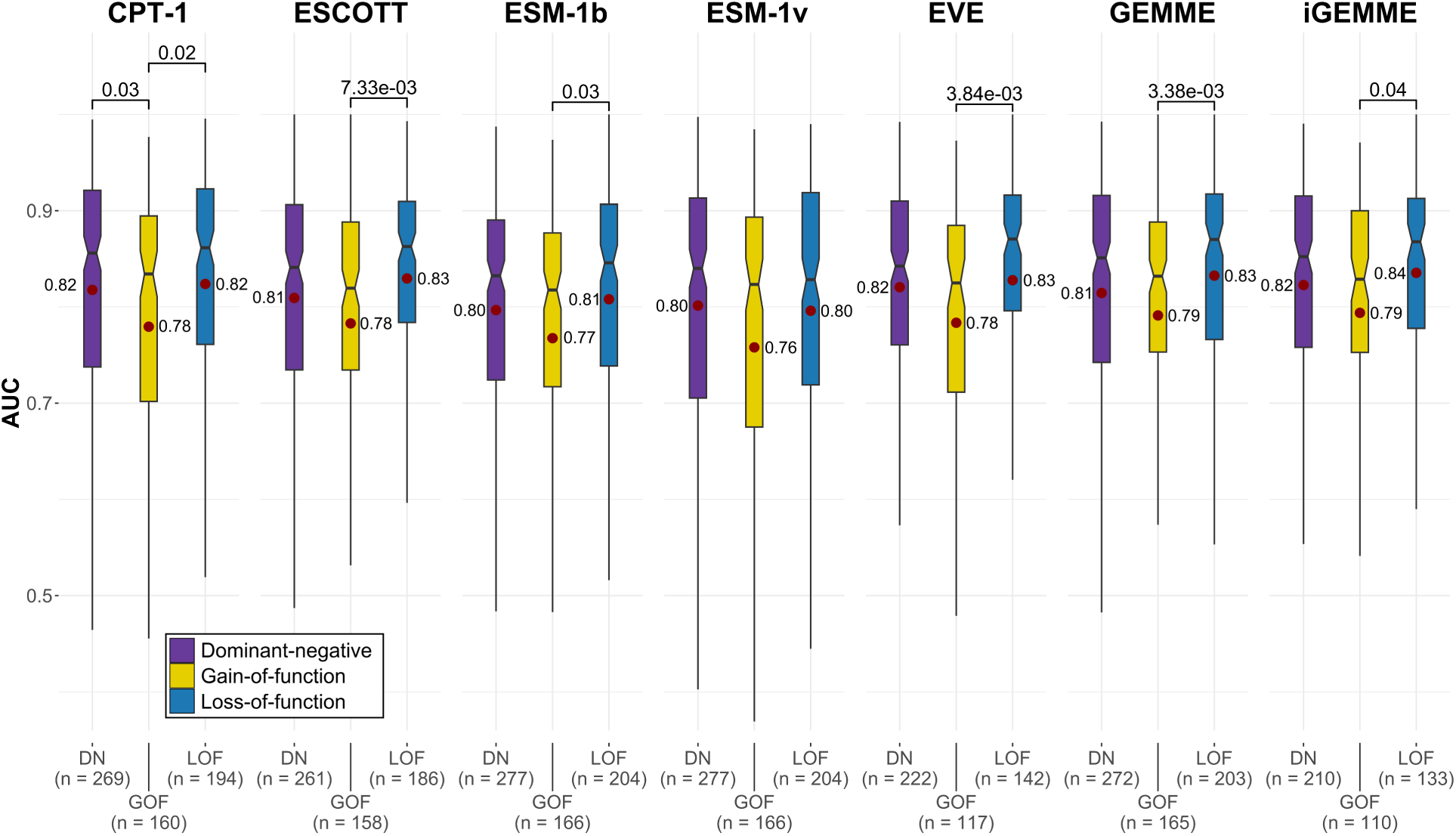
Top-performing population-free VEPs still show a degree of underperformance in GOF disease genes. Boxes denote data within 25th and 75th percentiles, and contain median (middle line) and mean (red dot) value notations. Whiskers extend from the box to furthest values within 1.5x the inter-quartile range.

**Supplementary Figure 5.**
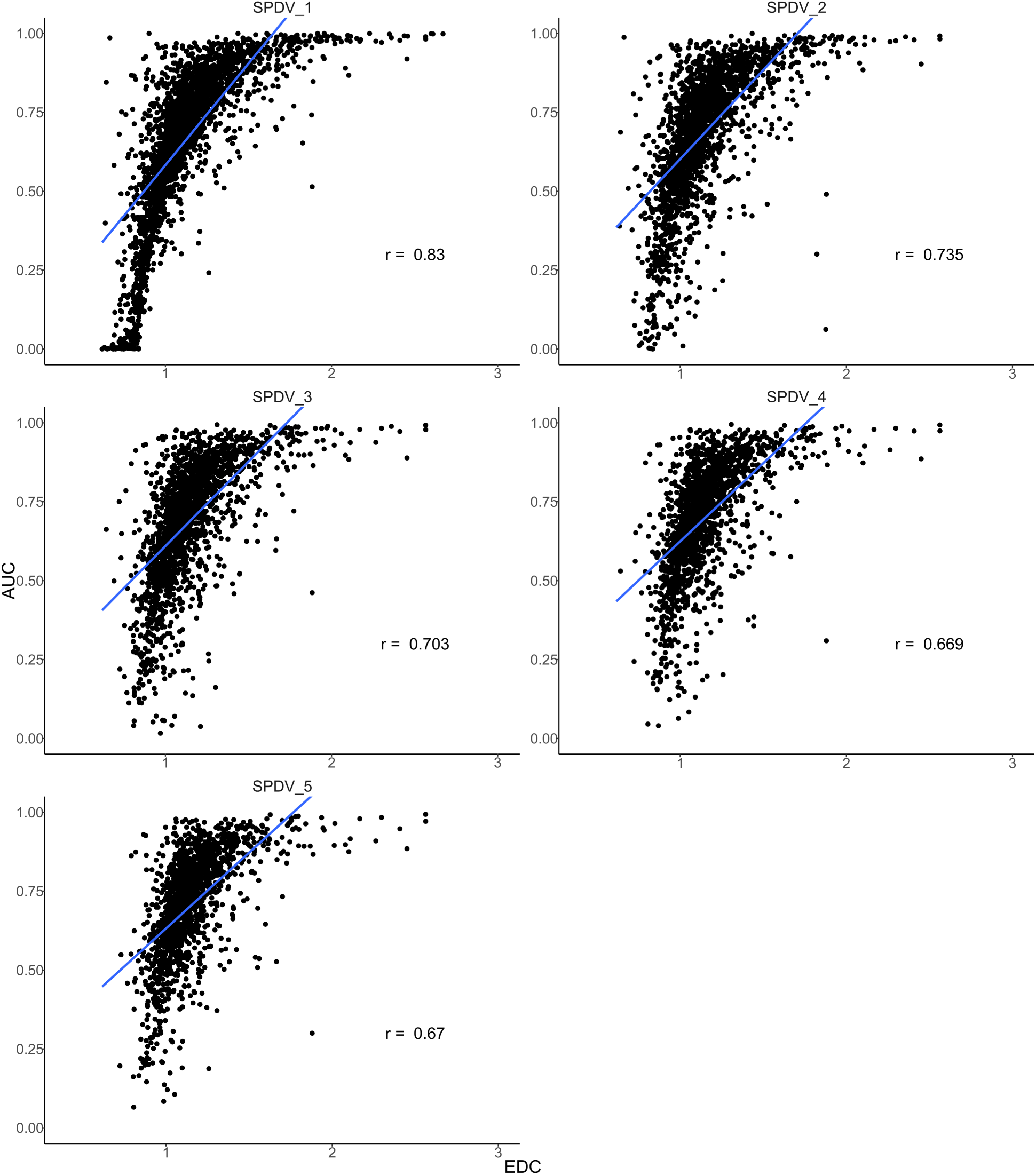
Per-gene EDC values are highly predictive of the SPDV metric performance. Rho value represents Spearman’s correlation.

**Supplementary Figure 6.**
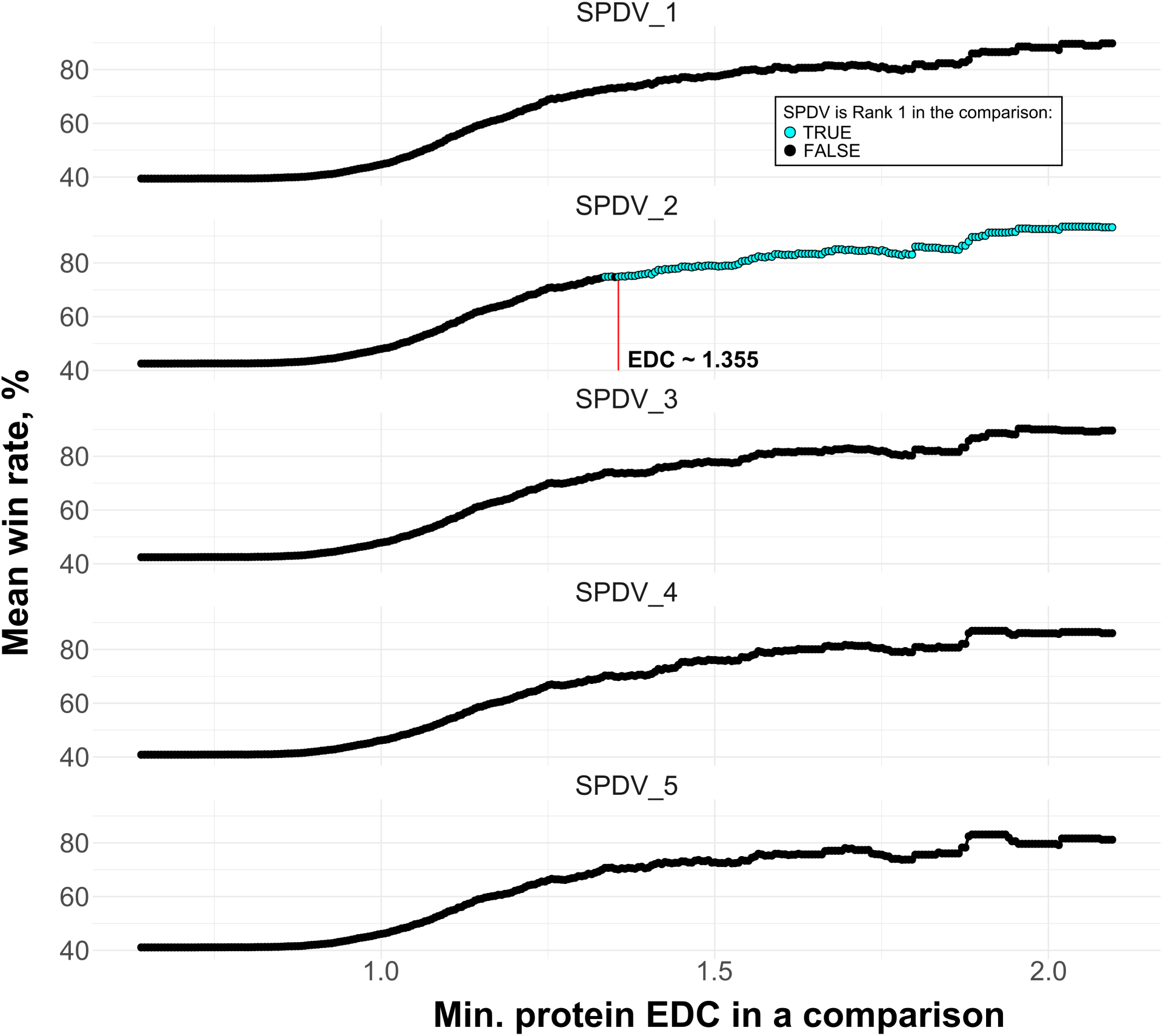
In proteins exceeding a certain degree of disease variant clustering, SPDV_2 consistently outperforms other VEPs and SPDVs in disease variant identification. The benchmark was carried out starting from a set of 1,875 disease genes with at least 5 distinct disease positions, annotated with EDC values. An EDC step of 0.05 was used to iteratively filter out genes to assess the performance of the five SPDV metrics and other population-free VEPs in a win rate competition. The red line marks the point of EDC ∼1.355 where SPDV_2 becomes the best overall predictor.

**Supplementary Figure 7.**
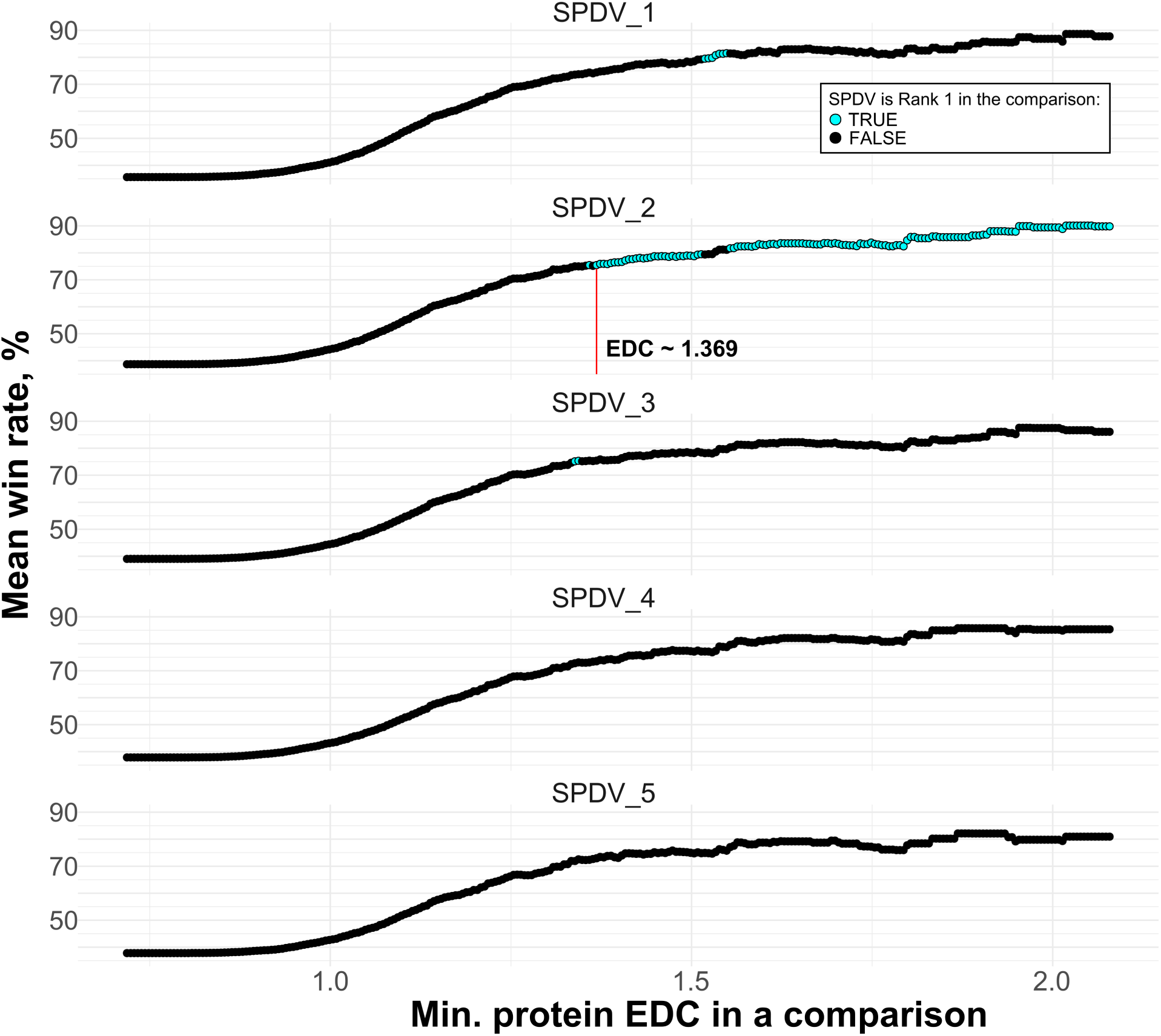
Removal of variants in disordered regions does not drastically impact the EDC threshold whereafter SPDVs become consistently the most accurate at identifying disease variants. The benchmark was carried on a new list of 1,726 genes that had at least 5 distinct disease positions in ordered (pLDDT>70) areas of the structure. An EDC step of 0.05 was used to iteratively filter out genes to assess the performance of the five SPDV metrics and other population-free VEPs in a win rate competition. The red line marks the point of EDC ∼1.355 where SPDV_2 becomes the best overall predictor.

**Supplementary Figure 8.**
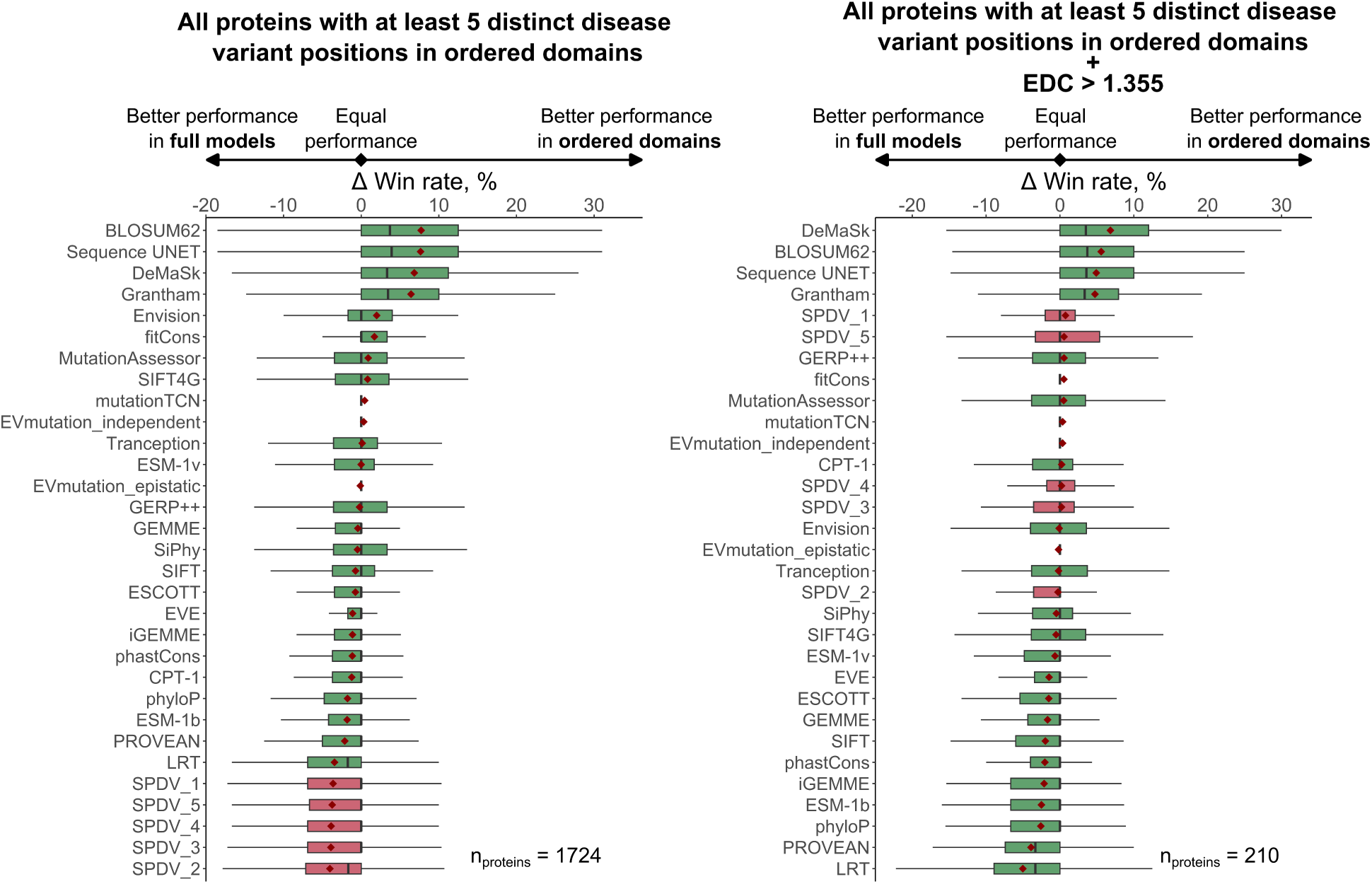
While the perceived SPDV performance is generally inflated due to easily distinguishable benign variants in disordered regions, the ‘low-hanging fruit’ variants do not influence SPDV performance in the highly-clustered disease variant gene set. Win rates were calculated using a pairwise AUC comparison scheme (see Methods). The difference in average win rates was derived on a per-gene basis, subtracting the win rate calculated for the full model from the one derived only using variants from structured residues (pLDDT>70). Boxes denote data within 25th and 75th percentiles, and contain median (middle line) and mean (red dot) value notations. Whiskers extend from the box to furthest values within 1.5x the inter-quartile range.

